# Mutation rate, selection, and epistasis inferred from RNA virus haplotypes via neural posterior estimation

**DOI:** 10.1101/2023.01.09.523230

**Authors:** Itamar Caspi, Moran Meir, Nadav Ben Nun, Uri Yakhini, Adi Stern, Yoav Ram

## Abstract

RNA viruses are particularly notorious for their high levels of genetic diversity, which is generated through the forces of mutation and natural selection. However, disentangling these two forces is a considerable challenge, and this may lead to widely divergent estimates of viral mutation rates, as well as difficulties in inferring fitness effects of mutations. Here, we develop, test, and apply an approach aimed at inferring the mutation rate and key parameters that govern natural selection, from haplotype sequences covering full length genomes of an evolving virus population. Our approach employs *neural posterior estimation*, a computational technique that applies simulation-based inference with neural networks to jointly infer multiple model parameters. We first tested our approach on synthetic data simulated using different mutation rates and selection parameters while accounting for sequencing errors. Reassuringly, the inferred parameter estimates were accurate and unbiased. We then applied our approach to haplotype sequencing data from a serial-passaging experiment with the MS2 bacteriophage. We estimated that the mutation rate of this phage is around 0.2 mutations per genome per replication cycle (95% highest density interval: 0.051-0.56). We validated this finding with two different approaches based on single-locus models that gave similar estimates but with much broader posterior distributions. Furthermore, we found evidence for reciprocal sign epistasis between four strongly beneficial mutations that all reside in an RNA stem-loop that controls the expression of the viral lysis protein, responsible for lysing host cells and viral egress. We surmise that there is a fine balance between over and under-expression of lysis that leads to this pattern of epistasis. To summarize, we have developed an approach for joint inference of the mutation rate and selection parameters from full haplotype data with sequencing errors, and used it to reveal features governing MS2 evolution.

## Introduction

Mutations are one of the primary sources of genetic heterogeneity in viruses and can be seen as the fuel of evolution. The mutation rate is defined as the number of new mutations in a genome over a unit of time, usually one generation. Viruses are notorious for their extremely high mutation rates (Duffy, 2018; Sanjuán et al., 2010; Zanini et al., 2017). Therefore, the viral mutation rate is a key parameter of virus evolution that, together with selection, determines the extent to which genetic diversity is created in a population of viruses.

Current methods for measuring the mutation rate often involve genomic sequencing of viral populations across time. Experimental evolution of viral populations is a powerful way to track viral mutations: in a controlled laboratory serial passaging experiment, a viral population is allowed to replicate for several generations, and deep sequencing is used to measure the dynamics of mutant allele frequencies. The high viral mutation rate will lead to many new mutations being constantly introduced; selection will eliminate deleterious mutations and will lead to an increase in frequency of beneficial mutations that promote viral fitness. The challenge is then, when observing mutations and their frequencies across time, to separate between the effects of mutation and natural selection (Peck & Lauring, 2018). One way of overcoming this challenge is to focus on specific mutations with known fitness effects. For example, one may focus on lethal mutations and measure their frequency, because under mutation-selection balance, the frequency of lethal mutations should be equal to the mutation rate (Acevedo et al., 2014; Cuevas et al., 2009; Sanjuán et al., 2010). Focusing on lethal mutations leads to several subsequent challenges: first, defining mutations as lethal is not always straightforward; second, those lethal mutations that can be defined may be a small subset of all loci; finally, lethal mutations may not be easily distinguished from sequencing errors (but see (Acevedo et al., 2014)).

Another approach is to focus on neutral mutations. In theory, if evolution begins with a completely homogenous population, neutral mutations are expected to accumulate at the rate of mutation. The challenge, then, is to specify which mutations are neutral. In some studies, synonymous mutations are assumed to be neutral (Stern et al., 2017; Tromas & Elena, 2010; Zanini et al., 2017; Zinger et al., 2019). However, there is growing evidence that many synonymous mutations are not neutral, and this may be particularly exacerbated in viruses with small dense genomes where genomic regions may have overlapping functions (Cuevas et al., 2012; Mayrose et al., 2013; Zanini & Neher, 2013). An additional complication is that both the lethal mutation and neutral mutation approaches are based on single-locus models, neglecting multi-locus effects such as background selection, selective sweeps (Feder et al., 2021), and epistatic interactions.

Here, we focus on MS2, +ssRNA virus from the Leviviridae family, which is a widely studied model virus. Nevertheless, the mutation rate of MS2 has yet to be estimated. Mostly, estimates of mutation rates of Qβ, a close relative of MS2, are assumed to apply for MS2. These estimates vary widely, ranging from 0.08 (García-Villada & Drake, 2012) to 0.6 (Bradwell et al., 2013) to 6.5 (Batschelet et al., 1976; Domingo et al., 1976; Drake, 1993) mutations per genome per replication cycle. This variation suggests that previous estimation approaches, based on only a handful of genomic loci, are not accurate enough (García-Villada & Drake, 2012). Importantly, MS2 has a particularly short genome of about 3,500 bases, allowing us to obtain reads that cover the entire genome.

Here, we develop and test an approach to jointly infer both the mutation rate and selection parameters from sequencing data of MS2. Our approach relies on key methodological novelties: (a) long-read sequencing that covers the full length of the viral genome (i.e., haplotypes) (Callahan et al., 2021); (b) a multi-locus evolutionary model that captures the multitude of segregating haplotypes in the population; and (c) deep artificial neural networks and simulation-based inference that allow efficient and high-dimensional inference of model parameters (Avecilla et al., 2021; Cranmer et al., 2020). We applied this approach to infer the MS2 mutation rate and selection parameters from long-read haplotype sequences sampled from populations evolving in the lab.

## Methods

### Evolutionary experiment

Clonal MS2 stock was propagated from a single plaque, to ensure that the experiment began with a phage population as genetically homogeneous as possible. Our experimental design was similar to the one described previously (Meir et al., 2020) with some changes: we performed ten serial passages at 37°C with three biological replicates (lines A-C). The serial passaging protocol was designed to allow tight population size control, to limit host–phage coevolution (naïve *Escherichia coli* c-3000 hosts were provided for each passage), and to limit co-infection and ecological interactions of phages within the cell, as described below.

We measured the length of the MS2 replication cycle by performing a onestep growth curve experiment as described in detail before (Meir et al., 2020). Briefly, after infection of the *E. coli* host cells we measured the number of MS2 particle forming units (PFU) per ml at various time points across 240 minutes. After 120 minutes we reached the maximal PFU of viruses after infection, and thus passages were arrested after 120 minutes.

The serial passages were performed at a multiplicity of infection (MOI) of 0.1 as follows: 10 ml cultures of naïve *E. coli* c-3000 were grown to an optical density of OD_600_=0.4 (corresponding to a density of about 2·10^7^ cells/ml). Each passage was infected with 0.2 ml of 10^8^ phages from the previous passage, thus keeping an MOI of 0.1 PFU per cell (*N*=2·10^7^ PFU). The cultures were grown for 120 minutes with shaking of 200 rpm, and the *E. coli* cells were then removed by centrifugation. The supernatant was subjected to filtration with a 0.22 μm filter to remove any remaining residues. The new phage stock was then stored at 4°C. Aliquots of these phage stocks were used for measuring the concentration of phages by plaque assay (as described below), infecting the next serial passage, isolating RNA for whole genome deep sequencing (as described below) and maintaining a frozen stock of the evolving lines in 15% glycerol at -80°C. RNA was isolated using the QIAamp® viral RNA Mini kit (QIAGEN) according to the manufacturer’s instructions.

### Plaque assay

Plaque assay was performed at the end of each passage to determine the MS2 phage titer (Meir et al., 2020). Briefly, *E. coli* c3000 was grown to mid-logarithmic phase (OD_600_ of 0.5) in rich growing medium LB at 37°C with shaking at 200 rpm. Serial dilutions of the MS2 samples were prepared in NaCl 0.85% to reduce the phage concentration to below 200 PFU/ml (could be counted easily on a Petri dish with the naked eye). We added into each test tube 5 ml of soft agar (70%) with 1 ml of *E. coli*. Then, 0.1 ml of each phage sample was added, and all of the tube content was emptied onto solid base agar in standard Petri dishes and allowed to harden. The plates were incubated overnight at 37°C.

### Loop genomics library construction, sequencing, and processing

RNA from passages 3, 7, and 10 from all three lines was sent to Loop Genomics (San Jose, CA, USA) and sequenced using the LoopSeq RNA preparation kit and its protocol (information available at loopgenomics.com). Loop Genomics’ synthetic long-read approach enabled full-length sequencing of the MS2 RNA genome with a low error rate of 10^−5^ errors per base (Callahan et al., 2021). In short, the LoopSeq method uses unique molecular barcoding labeling technology in which each barcode is distributed across a single genome followed by fragmentation of the genome to shorter fragments. The labeled fragmented genomes are then sequenced by existing standard short-read sequencing approaches on the Illumina sequencing platforms. The short-reads raw data was uploaded to the Loop Genomics’ unique analytic pipeline. The pipeline was used for low-quality base trimming, unique sample barcode demultiplexing, and synthetic long-read reconstruction. The synthetic long-read reconstruction is a process that enables the de novo assembly to the full-length genomes after rearranging the short reads tagged with the same unique barcode.

### Data analysis

We ran BLAST (Altschul et al., 1990; McGinnis & Madden, 2004) to align the long reads obtained from LoopSeq against the MS2 reference sequence (GenBank ID V00642.1, with some small differences noted in (Meir et al., 2020)) with parameters: --evalue 1e-07 --perc_identity 0.85 --task blastn --num_alignments 1000000 --dust no --soft_masking F. In order to ensure we obtained only reads that spanned the entire alignment, we then filtered out any alignments that aligned more than once to the reference, and filtered alignments that were shorter than 3,500 nucleotides (98% of the length of the reference genome). We further removed alignments that resulted in a minus strand alignment, i.e., were the reverse complement of the genome.

As described below, our model relies on single-nucleotide variation (SNV) counts, and in particular categorizes them as synonymous or non-synonymous. We thus excluded both the untranslated regions of the genomes (which are very short and account for less than 10% of the genome), as well as indels from our analysis, as their biological effects are more complex to analyze.

### Evolutionary model

Instead of considering each SNV in the genome, we label each SNV as belonging into one of the following types based on whether a mutation at a site is: (i) synonymous or not; (ii) beneficial or non-beneficial, and (iii) whether it was part of the founding population or not. We group genotypes together by the number of each of these mutation types into genotype classes (illustrated in Fig. 1). Monitoring just the genotype classes, rather than the genotypes themselves, reduces the number of model parameters to only ten and makes our model computationally tractable.

**Figure 1.**
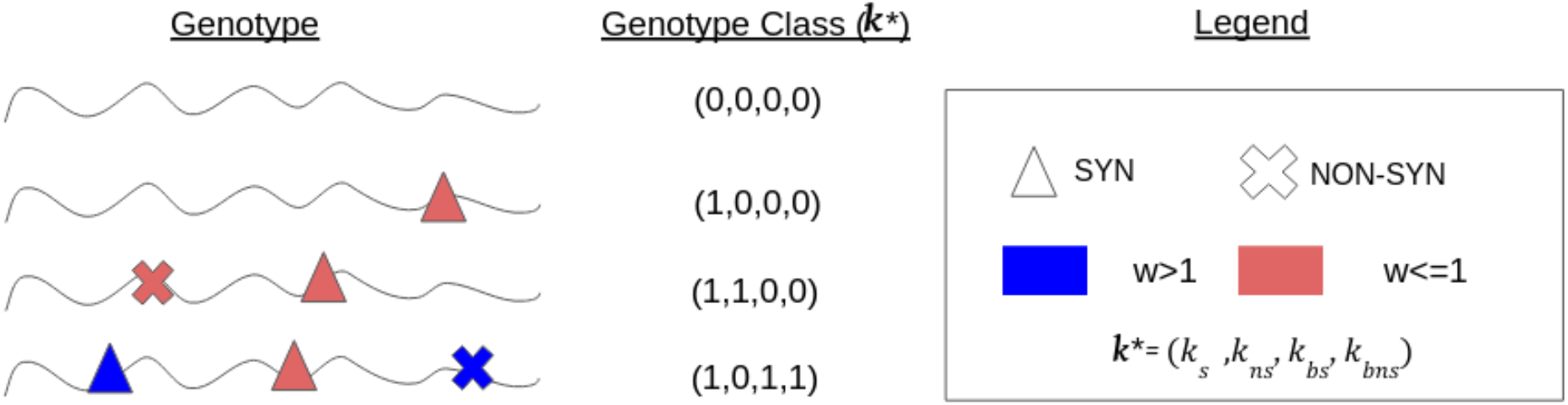
Illustration of genotype Classes. The basis of our method relies on assigning genotypes to genotype classes. Each genotype class *k* is a 6-tuple that represents all genomes with the same number of non-beneficial synonymous SNVs that derive from the initial population, non-beneficial non-synonymous SNVs from the initial population, non-beneficial synonymous SNVs, non-beneficial non-synonymous SNVs, beneficial synonymous SNVs, and beneficial non-synonymous SNVs. For simplicity, simplified genotype classes (*k\**) consisting of only 4-tuples representing the last four types of mutations (i.e., excluding the initial founder population SNVs) are represented in the figure. SNVs of the same type are interchangeable and are assumed to have the same fitness effect (*w*). Genotype classes allow our model to be computationally tractable, while still giving a detailed account of the evolutionary dynamics.

#### Initial founding population

Our experimental conditions were designed to ensure a homogeneous initial population, derived from a plaque. However, the generation of this initial population still included multiple replication cycles at slightly different conditions than our experiment, such as higher MOI. We thus directly modeled the limited diversity present in the initial population. We assumed that the average number of synonymous SNVs per genotype in the initial population is *M*_*s*_ and the average number of non-synonymous SNVs per genotype in the initial population is *M*_*ns*_. We assume the number of SNVs in the initial population is *M*∼*Poi*(*M*_*s*_ + *M*_*ns*_) and for simplicity also assumed that this plaque-derived founding population does not contain beneficial SNVs. Therefore, the number of synonymous *k*_*Is*_ and non-synonymous *k*_*Ins*_ mutations per individual in the initial population are

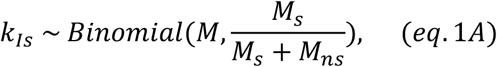

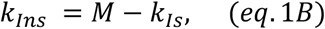

#### Evolution during serial passaging

We model the evolutionary dynamics using a Wright-Fisher framework with constant population size *N* and non-overlapping generations. We follow the frequency of genotypes with exactly *k*_*Is*_ non-beneficial synonymous SNVs that derive from the initial population, *k*_*Ins*_ non-beneficial non-synonymous SNVs from the initial population, *k*_*s*_ non-beneficial synonymous SNVs, *k*_*ns*_ non-beneficial non-synonymous SNVs, *k*_*bs*_ beneficial synonymous SNVs, and *k*_*bns*_ beneficial non-synonymous SNVs, such that each genotype can be classified into a class *k = (k*_*Is*_, *k*_*Ins*_, *k*_*s*_, *k*_*ns*_, *k*_*bs*_, *k*_*bns*_), *k*_*i*_ ≥ 0.

##### Mutation

We assume the number of new SNVs per genotype per replication cycle is Poisson distributed with expected value *U*, which is defined as the mutation rate in mutations per genome per replication cycle (Tenaillon et al., 1999). The probability that a SNV is synonymous or nonsynonymous is given by *p*_*s*_ and *p*_*ns*_ = 1 − *p*_*s*_, respectively. The probability that a synonymous or nonsynonymous SNV is beneficial is *p*_*bs*_ and *p*_*bns*_, respectively. Thus, the number of new SNVs per genotype is given by

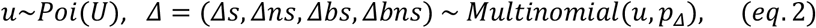

Where *u* is the total number of new SNVs in one individual, *Δi* is the number of new SNVs of type *i*, and *p*_*Δ*_ = (*p*_*s*_ − *p*_*bs*_, *p*_*ns*_ − *p*_*bns*_, *p*_*bs*_, *p*_*bns*_) (which sums to 1). Thus, the change in *f*_*k*_, frequency of genotype class *k*, due to mutation is given by

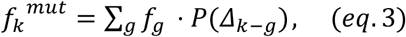

where *g* is an index over all existing genotype classes that can mutate to genotype class *k*; *Δ*_*k*−*g*_ is the differences in the number of mutations of the various types between genotype class *k* and *g*; and *P(Δ*_*k*−*g*_) is the Poisson-multinomial probability mass function (*eq*. 2). Since the poisson-multinomial distribution has infinite support, we truncate the distribution by setting *P*(*Δ*_*k*−*g*_) = 0 if *P*(*Δ*_*k*−*g*_) < 1/(100*N*). We neglect the effect of recombination due to low MOI.

##### Fitness

For simplicity, we assume a single value for the fitness effect of synonymous non-beneficial SNVs, *w*_*s*_; and a single value for fitness effect of non-synonymous non-beneficial SNVs, *w*_*ns*_.

Beneficial SNVs (synonymous or non-synonymous) are modeled separately, and the fitness effect of beneficial SNVs (synonymous or non-synonymous) is *w*_*b*_.

##### Initial founder population fitness

Since the initial population evolved under possibly different selection pressures (described above), we allow SNVs in the initial population to have different fitness effects, but we assume the log-fitness of these initial SNVs are correlated with the log-fitness of non-beneficial SNVs in the experiment, with correlation coefficient *δ*: here, *δ* = 0 implies the initial population SNVs are neutral under the experimental conditions; *δ* = 1 implies that the initial population SNVs fitness effects are the same as that of the non-beneficial SNVs; and *δ* > 1 implies the SNVs in the initial population are more deleterious than later non-beneficial SNVs.

#### Epistasis

We also model the potential interaction of multiple *beneficial* SNVs by introducing an epistasis parameter *η*: when *η* > 1 there is positive epistasis, when 0.5 < *η* < 1 there is negative epistasis; when *η* < 0.5 there is sign epistasis; and when *η* < 0 there is reciprocal sign epistasis so that the effect of two beneficial mutations is deleterious.

##### Fitness of genotype classes

We assume multiplicative fitness. Therefore, the fitness of genotype class *k=*(*k*_*Is*_, *k*_*Ins*_, *k*_*s*_, *k*_*ns*_, *k*_*bs*_, *k*_*bns*_) is

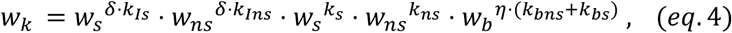

where *η =1* if *k*_*bns*_ + *k*_*bs*_ < 2.

Thus, the effect of natural selection on genotype frequencies is given by

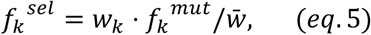

where 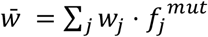 is the population mean fitness.

##### Random sampling

Serial passaging that includes sampling progeny virus and initiating the nest serial passage is modeled by assuming random sampling. Thus, the number of virions of each genotype class after sampling is given by *n* = {*n*_*k*_}∼*Multinomial*(*N, f*^*sel*^), where *f*^*sel*^ = {*f*_*k*_^*sel*^} and *k* is an index over all existing genotype classes. Accordingly, the frequency of genotype class *k* in the next generation is

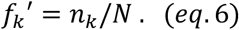

We simulate the stochastic genotype frequency dynamics in the experiment by iterating eq 1-6 *f*_*k*_ for multiple generations.

##### Sequencing process

Because only a sample of genomes are sequenced, and sequencing is error prone, we model sequencing errors and sampling. The frequency 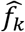 of genotype class *k* after sampling and sequencing is

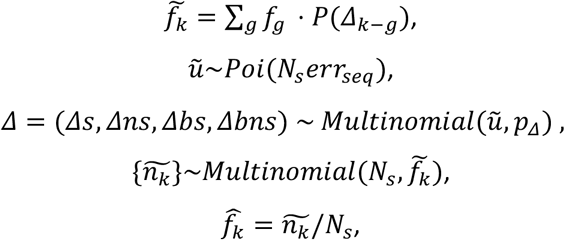

where *N*_*s*_ is the sample sequencing coverage (see below) and *err*_*seq*_ is the expected average sequencing error set at 10^−5^ errors per base, based on the reported LoopSeq error rate (Callahan et al., 2021). We further tested *err*_*seq*_= 10^−4^ errors per base to assess the robustness of our method to a higher error rate.

#### Experimental parameter values

A viral population size *N* = 2 · 10^8^ was assumed corresponding to the experimental setup, see above. Each line and passage had different sequencing coverage *N*_*s*_ (mean = 2,589, standard deviation = 1,195, minimum = 1,309, maximum = 4,976; see Table S1 for details). The probability that a SNV is synonymous is *p*_*s*_ = 0.28, corresponding to the possible point synonymous and nonsynonymous SNVs in the MS2 reference genome. Other parameter values were inferred from experimental data, see next section.

### Bayesian inference

#### Summary statistics

We applied three summary statistics: short-reads summary statistic (SR), long-reads summary statistic (LR), and labeled long-reads summary statistic (L-LR). The simplest, SR, is a vector of the average number of synonymous and nonsynonymous SNVs per genotype at each passage *t*. Thus, SR is a vector with two entries per passage and six entries in total.

Using LoopSeq long-read sequencing, which covers the entire MS2 genome, we could use a more informative summary statistic, LR, which counts the frequency of genotypes containing exactly *k*_*Is*_ + *k*_*s*_ + *k*_*bs*_ synonymous SNVs and *k*_*Ins*_ + *k*_*ns*_ + *k*_*bns*_ nonsynonymous SNVs respectively, at passage *t*. We counted up to ten SNVs of each type, producing 66 possible SNV combinations per passage. We also included SR in LR, producing a vector with 68 entries per passage and 204 entries in total.

We used a similar but more informative summary statistic, L-LR, which includes the number of beneficial SNVs. We labeled four SNVs as beneficial: they were all rare at passage 3 but reached a frequency higher than 3% in all three lines by passage 10 (Fig. 2A, purple lines). Assuming that the per base mutation rate is at most 0.001 (Drake, 1993), a neutral SNV is expected to be at a frequency of about 1% by passage 10. Labeling the beneficial SNVs allowed us to obtain the frequencies of genotypes containing exactly *k*_*Is*_ + *k*_*s*_ synonymous non-beneficial SNVs, *k*_*bs*_ synonymous beneficial SNVs, *k*_*Ins*_ + *k*_*ns*_nonsynonymous non-beneficial SNVs, and *k*_*bns*_ nonsynonymous beneficial SNVs at each passage *t*. We counted up to ten SNVs of each of the four types, producing 1,001 frequency values per passage. Including SR in L-LR produced a vector with 3,009 entries in total.

**Figure 2.**
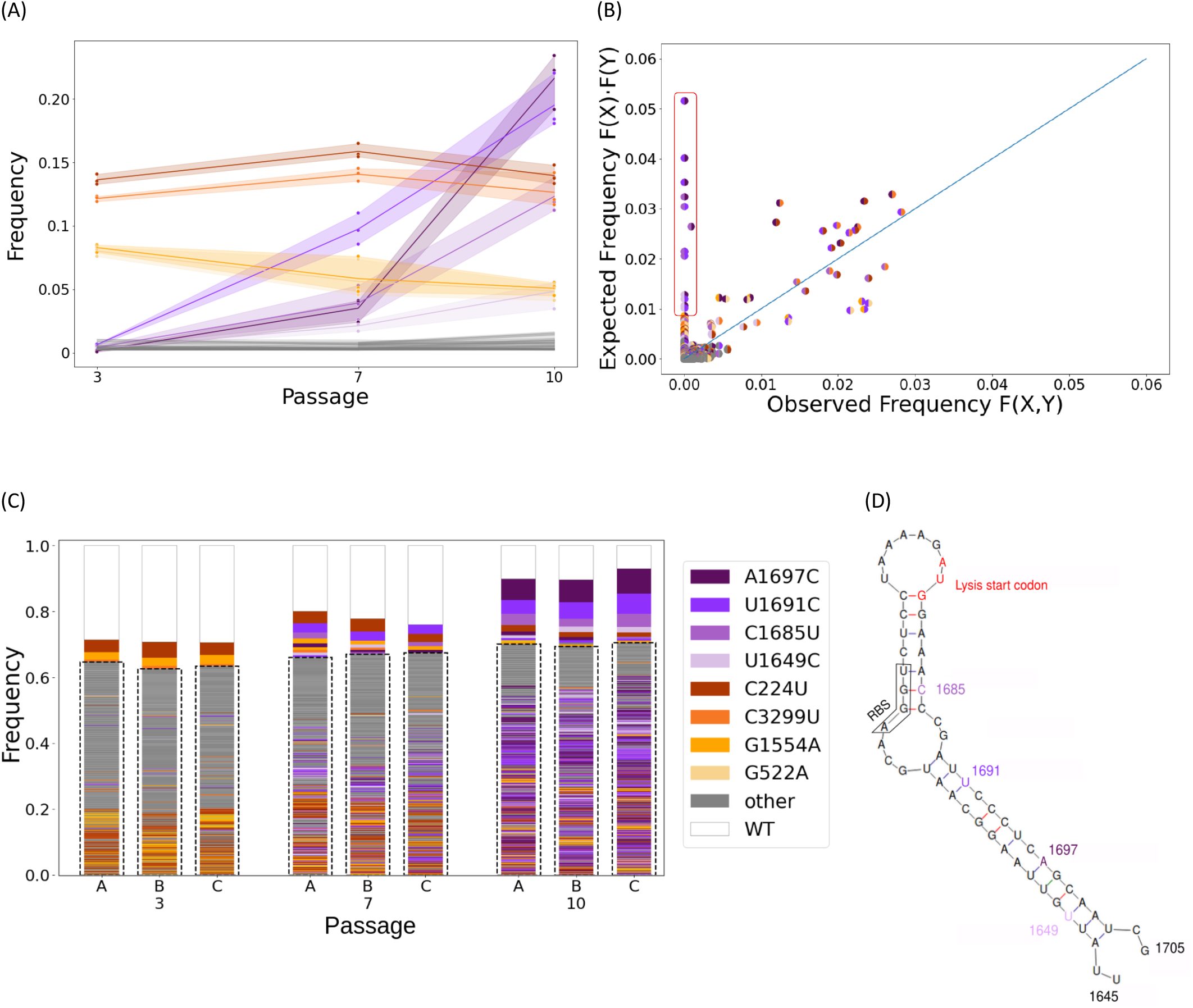
SNV and genome frequencies across time. **(A)** Markers show SNV frequencies in the three replicas, with high-frequency SNVs in purple, founder SNVs in brown-orange, and all other low-frequency SNVs in gray. The area between the replicas is shaded. **(B)** Observed versus expected frequencies of pairs of SNVs. For a pair of SNVs denoted by X and Y, *F(X)* and *F(Y)* are their frequencies across all genotypes, respectively; *F(X)F(Y)* is the expected frequency of the pair assuming independence; and *F(X,Y)* is the observed frequency of the pair across all genotype; frequencies are taken from all three replicas at passage 10. The diagonal line represents independence of the pair of SNVs, i.e., *F(X,Y)=F(X)F(Y)*. Founder SNV pairs excluded from this panel. Pairs of beneficial SNVs are surrounded by a red box. **(C)** Genetic diversity for each replica and passage. When a genome contains more than one high-frequency SNV, the color of its box is defined by the first SNV in the order of appearance in the legend. The genomes are ordered by their frequency in each dataset and the number of SNVs within them. Rare genotypes are defined as present in less than 0.5% of a sample and are boxed with a dashed line. The RNA structure where all beneficial SNVs occur, inferred using *Mfold* (Zuker, 2003) and in line the experimentally resolved structure (Dai et al., 2017). This region spans the end of the coat gene and the overlapping lysis gene. All beneficial SNVs are synonymous with respect to the coat gene and nonsynonymous with respect to the lysis gene. RBS marks the ribosomal binding site for the lysis gene.

#### Prior distributions

We assumed uniform (noninformative) prior distributions with ranges set based on current estimates in the literature (Table 1). The prior distribution of *w*_*b*_ is much wider than the estimates in the literature to account for the rapid increase in frequencies of some SNVs we observe in the empirical data (Fig. 4B). Prior distributions for *w*_*s*_ and *w*_*ns*_ are wide to avoid bias in the posterior. Prior distributions of *M*_*s*_ and *M*_*ns*_ are based on experimental sequencing data from three different MS2 plaque sequences (data not shown).

**Table 1.** Model parameters.

| Description | Symbol | Prior distribution |
| --- | --- | --- |
| *average non-beneficial synonymous fitness effect* | $w_s$ | uniform(0.1, 1) |
| *average non-beneficial nonsynonymous fitness effect* | $w_{ns}$ | uniform(0.1, 1) |
| *average beneficial fitness effect* | $w_b$ | uniform(1, 3) (Sanjuán et al., 2004) |
| *beneficial synonymous SNV probability* | $p_{bs}$ | uniform(0, 0.01) (Sanjuán et al., 2004) |
| *beneficial nonsynonymous SNV probability* | $p_{bns}$ | uniform(0, 0.01) (Sanjuán et al., 2004) |
| *average number of synonymous SNVs per genotype in the initial population* | $M_s$ | uniform(0.4, 0.6) |
| *average number of nonsynonymous SNVs per genotype in the initial population* | $M_{ns}$ | uniform(0.7, 0.9) |
| *initial log-fitness correlation* | $\delta$ | uniform(0, 2) |
| *epistasis* | $\eta$ | uniform(-1, 3) |
| *genome-wide mutation rate* | $U$ | log-uniform(-4, 0.3)<br>(Sanjuán et al., 2010) |

#### Sequential neural posterior estimation (SNPE)

We used a recently developed neural-network-assisted likelihood-free inference method, *sequential neural posterior estimation* (SNPE) (Greenberg et al., 2019), or specifically, the SNPE-C implementation in the Python package *sbi* (Tejero-Cantero et al., 2020). SNPE has been recently applied for inferring the formation rate and fitness effect of copy number variation in populations of yeast evolving under nutrient limitation in a chemostat (Avecilla et al., 2021). Briefly, SNPE trains an artificial neural network on a training set of parameters (generated from the prior distribution) and simulated data (generated from the evolutionary model) to estimate the joint density of model parameters and data (conditioned on the prior distribution). This joint density is effectively an *amortized posterior distribution* of the model parameters, which can be evaluated for specific observed data. Amortization allows the evaluation of the posterior distribution for each experimental observation (i.e., replicate or line) without needing to re-train the neural network (in contrast to MCMC-based approaches that require a new run of the algorithm to infer a posterior from a new observation). As the neural density estimator, we used a *masked autoregressive flow* (MAF) (Papamakarios et al., 2018), which uses a stack of autoregressive models (specifically, *masked autoencoders for distribution estimation*, or MADE (Germain et al., 2015)) to model a sequence of transformations on random variables.

##### Ensemble SNPE

We further extend the SNPE method to the *ensemble SNPE* by averaging the posterior distribution estimated by *eight* density estimators, each independently trained with non-overlapping training sets of 10,000 simulations sampled from the same prior. Below, we compare the performance of the ensemble SNPE to *individual SNPE*, which we trained on a training set of 80,000 simulations.

#### Approximate Bayesian computation with rejection sampling (REJ-ABC)

We compared SNPE with a classical likelihood-free inference method, *approximate Bayesian computation with rejection sampling* (REJ-ABC) (Pritchard et al., 1999; Tavaré et al., 1997). ABC is used to approximate posterior distributions when an underlying likelihood function is analytically intractable. In REJ-ABC, parameter sets are independently sampled from the prior; a simulation is run for each parameter set; the distance between simulation results and observed data are computed; parameter sets that produced the lowest *ε*-percentile distances are accepted (while the rest are rejected); the posterior distribution is estimated from the accepted parameter sets. We used 80,000 parameter sets and simulations (the same that we used for SNPE) to produce the ABC-REJ posterior estimation. As a distance function, we used the root mean square error (RMSE) between the simulated and observed summary statistics. We produced posterior distributions using difference acceptance rates of *ε*=0.1%, 1% and 5%.

#### Parameter estimates and posterior predictive checks

To obtain a marginal maximum a-posteriori (MAP) estimate, we sample from the posterior distribution and construct a histogram with 100 bins. We then estimate the parameter as the center of the most frequent bin. We calculated highest density intervals (HDIs) using the Python package *ArviZ* (Kumar et al., 2019). We performed posterior predictive checks by simulating synthetic data with parameters sampled from the posterior and comparing it to the observed data.

#### Flexible Inference from Time-Series (FITS)

We also compared SNPE to a previous method developed by some of us called *FITS* (Zinger et al., 2019). In contrast to the above ABC-REJ and SNPE, FITS applies a single-locus Wright-Fisher model to all synonymous SNVs, assuming they are all neutral, and uses rejection sampling to approximate the posterior distribution of model parameters. Thus, FITS does not use the evolutionary model, prior distributions, and summary statistics described above.

#### Premature stop codons

We counted the number of premature stop codons across the genome in all reads from all passages and replicas and divided by the number of possible SNVs that could have resulted in a premature stop codon. Assuming premature stop codons are lethal, under a mutation-selection balance this ratio is expected to be equal to the mutation rate. The main problem with this approach is that observed premature stop codons due to mutation cannot be distinguished from sequencing errors (with a rate estimated at 0.005%, see above) and so this method only provides an upper bound on the estimated mutation rate.

### Data and code availability

Source code available at https://github.com/Stern-Lab/ms2-mutation-rate. Sequencing data available at https://www.ncbi.nlm.nih.gov/sra/PRJNA902661. Additional data available at https://zenodo.org/record/7486851.

## Results

### Individual SNV frequencies suggest positive selection

We began by inspecting the haplotype sequencing data, which was derived from passages 3, 7, and 10 from three independent biological replicas (A, B, and C). We first examined the SNV frequencies, focusing on those segregating at frequencies above 3% in all three replicas by the end of the experiment. Four SNVs were at a frequency above 5% already at passage 3, and remained at more or less constant frequencies throughout the experiment. Hence, we assumed they reflect standing variation in the initial founding population (Methods), and denote them as *founder SNVs* (Fig. 2A).

Conversely, we noted a set of four SNVs that increased dramatically in frequency, rising from less than 1% to frequencies ranging between ∼5% and ∼20% in all three replicas (Fig. 2A). This increase suggests that these four SNVs are under positive selection, and we thus denote them as *beneficial SNVs*.

We next examined haplotype composition, starting with pairs of SNVs. We compared the observed frequency of each SNV pair with its expected frequency, which is the product of their individual frequencies, assuming they are pairwise independent (Fig. 2B). We find that the four founder SNVs appear almost exclusively in two specific pairs (C3299T with C224T, and G1554A with G522A, Fig. 2A, not shown in 2B). In contrast, the four beneficial SNVs rarely or never appear on the same genome (one pair appears in one genome), exhibiting negative linkage disequilibrium. The low observed frequency of the beneficial SNV pairs can have two explanations: negative or sign epistasis between the beneficial SNVs, or a combination of strong selection and low mutation rate, which together do not allow enough time for accumulation of two beneficial SNVs on the same genome.

Next, we examined the genome diversity across replicas and passages. Given the data above, we noted that by passage 10, replicas A, B, and C respectively had 55%, 54% and 65% genomes with a beneficial SNV. We also observed high genome diversity across all replicas and all passages (Fig. 2C). Indeed, with the exception of the wildtype genome, which was present in 10%, 10%, and 7% in replicas A, B, and C, respectively, no genotype exceeded 7%, and 70% of genotypes were rare, i.e., present at frequencies lower than 0.5%. For example, the beneficial SNV A1697C reached a frequency of ∼20% by passage 10, but there was no single genotype bearing A1697C at a frequency above 7% (Fig. S6). This high level of diversity, including the abundance of rare genotypes, is suggestive of a high mutation rate, the occurrence of soft selective sweeps, and possible epistatic effects. We therefore developed a framework to jointly estimate the various evolutionary parameters from our time-series haplotype data (Methods).

### Validation of inference method on synthetic data

To validate the performance of our method, we simulated 2,000 synthetic data sets using the evolutionary model and a set of known parameter values (denoted as the “true” parameter values). We tested three summary statistics, each with subsequently more information than the previous one (SR, LR, and L-LR; see Methods).

#### Coverage

We first measured the *coverage*, defined as the proportion of inferences where the credible interval for a parameter contains the true parameter value (Prangle et al., 2014). We found that *individual SNPE* (a single neural density estimator trained on 80,000 simulations) produced posteriors that were overconfident, i.e., the 95% HDI contained the true parameter in less than 95% of the test cases (Fig. S1).

Indeed, it has been suggested that simulation-based inference methods may produce overconfident posterior approximations, and that this issue can be mitigated by using ensembles of estimators (Hermans et al., 2021). We therefore extended the individual SNPE to an *ensemble SNPE* that comprises *eight* individual SNPEs, each independently trained with non-overlapping training set (with a total simulation budget fixed at 80,000 simulations). We found that while individual SNPE mostly produces overconfident posteriors, the ensemble SNPE produces posteriors that are less confident, i.e., has superior coverage, compared to the individual SNPE (Fig. S1).

#### Parameter estimate accuracy

We compared our MAP estimates, which are point estimates of model parameters, with those of a simpler method, *approximate Bayesian computation with rejection sampling* (REJ-ABC). The latter was run with a 1% acceptance rate using the same data set of 80,000 simulations used to train the ensemble density estimators (Methods). REJ-ABC produced the largest MAP errors, on average, and did not seem to improve with more informative summary statistics (Fig. S2). For the SNPE inference methods, we found that more informative summary statistics produced similar or smaller MAP errors (Fig. S2). Ensemble SNPE performed similarly to the individual SNPE (Fig. S2). We chose to henceforth focus on ensemble SNPE because it had better coverage in most cases, and especially for the mutation rate (Fig. S1).

As expected, the more informative the summary statistic, the narrower the posterior distributions, and thus, the higher the information gain calculated by the Kullback-Leibler divergence (Fig. S3). The MAP error ratios of the ensemble SNPE were centered around zero, suggesting they were unbiased. We also observed a slight improvement in the MAP error ratio when increasing the information content of the summary statistics. All ensemble SNPE produced lower absolute MAP error ratios compared with REJ-ABC (Fig. 3).

**Figure 3.**
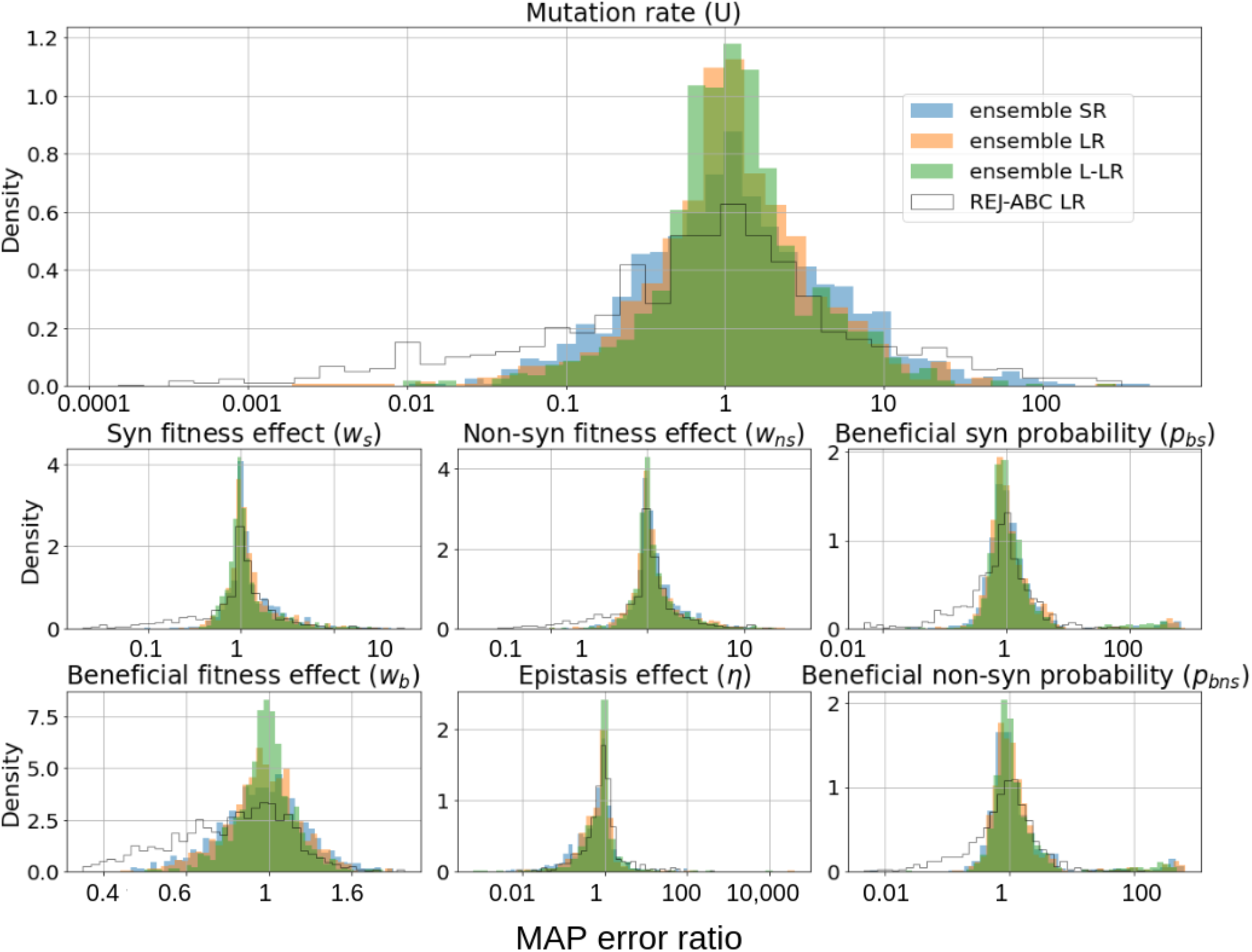
Parameters estimation accuracy on synthetic data. Log ratio of maximum a-posteriori (MAP) estimate and true parameter value on 2,000 synthetic datasets. Inferences used SNPE with three different summary statistics (short-reads: SR, blue; long-reads: LR, orange; labeled long-reads: L-LR, green) and REJ-ABC (LR, white).

### Inference of evolutionary parameters from empirical data

#### Mutation rate

We next applied the ensemble SNPE to the haplotype data from the MS2 experiment (see Fig. 2A), averaging the estimated posteriors over the three experimental replicas, and assuming sequencing errors at a rate of 5·10^−5^ per base, as reported by *Loop Genomics* (Callahan et al., 2021). We found that the ensemble SNPE with the SR and LR summary statistics produce similar posteriors, whereas using the L-LR summary statistic produced a narrower posterior distribution, and reproduced the dynamics of the evolving population (Fig. S8). All MAP estimates of the mutation rate *U* are between 0.15-0.2 mutations per genome per replication cycle (Table 2) and the widest HDI 95% is between 0.016 and 1.6. Given a genome length of 3,569 bases, this corresponds to between 4.2·10^−5^ and 5.6·10^−5^ SNVs per base per replication cycle.

**Table 2.** Mutation rate estimates. Maximum a-posteriori (MAP) estimates and 95% highest density intervals (HDI) of the posterior distributions of the genomic mutation rate U from Fig. 4A. Note that all estimates are similar, but ensemble SNPE produced HDI that is two orders of magnitude narrower than FITS (single-locus Wright-Fisher model with rejection sampling applied to all synonymous SNVs).

| Method | MAP estimate of $U$ | HDI 95% |
| --- | --- | --- |
| FITS | 0.197 | (0.0004, 0.796) |
| Premature stop codon frequency* | $\leq 0.187$ | N/A |
| Ensemble SNPE<br>Short-reads summary statistic (SR) | 0.161 | (0.013, 1.396) |
| Ensemble SNPE<br>Long-reads summary statistic (SR) | 0.221 | (0.014, 1.548) |
| Ensemble SNPE<br>Labeled long-reads summary statistic (L-LR) | 0.194 | (0.056, 0.73) |

\* Frequency of premature stop codons does not produce an HDI; rather, it produces an upper bound on the mutation rate and therefore reflects either the mutation rate or the sequencing error.

#### Comparison to other methods

We next compared our ensemble SNPE inference method to two alternative mutation rate inference methods: one based on the frequency of premature stop codons, and the other, FITS (Zinger et al., 2019), which is based on changes in the frequency of neutral SNVs (see Methods). We found that the MAP estimates from SNPE are in line with the frequency of premature stop codons, which is highly susceptible to sequencing errors and is therefore only an upper bound. Furthermore, SNPE produces a substantially narrower posterior distribution compared to FITS, regardless of the summary statistic (Fig. 4A, Table 2).

**Figure 4.**
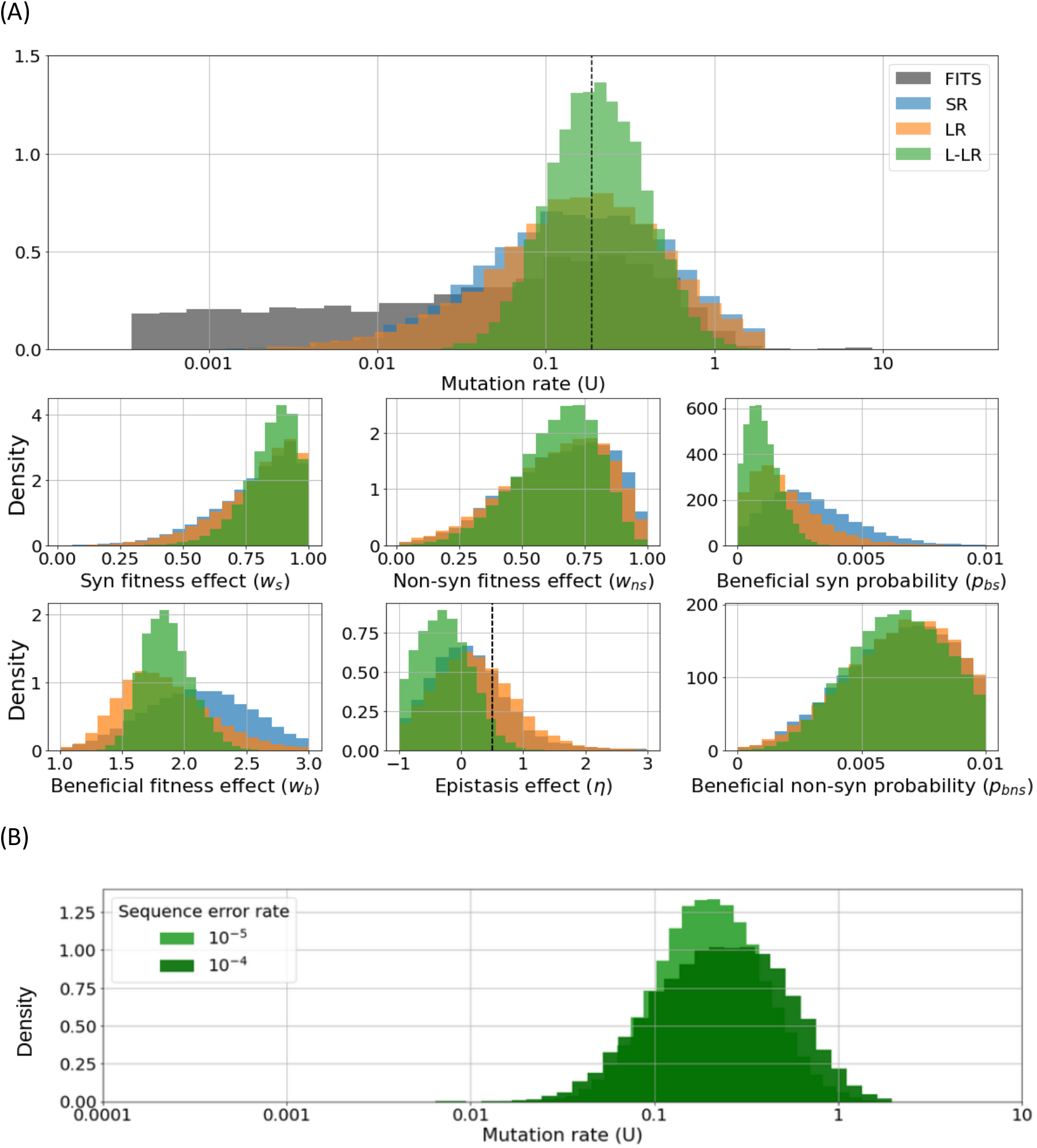
Posterior distributions of model parameters inferred for MS2. **(A)** Posterior distributions of the mutation rate using three summary statistics for the mutation rate produce a substantially narrower distribution compared to FITS (single-locus Wright-Fisher model with rejection sampling applied to all synonymous SNVs). The premature stop codon estimate is marked by a dashed line and is an upper bound on the mutation rate. Marginal posteriors of model parameters using ensemble SNPE with three different summary statistics. In most cases, the labeled long-reads summary statistic (L-LR) produces a narrower posterior and generally all posteriors agree. For epistasis, left of the dotted line are values indicating sign epistasis. **(B)** Marginal posterior distribution of the mutation rate *U* inferred with ensemble SNPE with the L-LR summary statistic compared to an estimate that assumes a tenfold increase in the sequencing error rate.

#### Sequencing error rate

Reassuringly, when assuming a tenfold increase in the sequence error rate (5·10^−4^) inference with the L-LR summary statistic was robust, i.e., the posterior distribution of the mutation rate and most other parameters remained similar (Fig. 4C, Fig. S9), although the mode of the posterior distributions of non-synonymous fitness effect (*w*_*ns*_) and beneficial non-synonymous probability (*p*_*bns*_) shifted under a higher error rate. However, using the SR and LR summary statistics produced wider posterior distributions for most model parameters (Figs. S4-S5). These results underscore the inherent difficulty in inferring mutation rates and fitness effects with error-prone short-read sequencing approaches and suggest that there is much added value in labeling beneficial SNVs when possible.

#### Fitness effects

We report estimates from inference with the ensemble SNPE with the L-LR summary statistic given its superior performance (Table S2). We estimate the non-beneficial synonymous mutation fitness effect *w*_*s*_ to be about 0.9 (0.651-1, HDI 95%; Table S2), implying that most synonymous mutations are neutral or slightly deleterious (Cuevas et al., 2012; Sanjuán et al., 2004). The estimated non-beneficial nonsynonymous mutation fitness effect is about 0.7 (0.311-0.922 HDI 95%; Fig. 4B; Table S2), this implicitly averages slightly deleterious, deleterious, and lethal mutations. The fitness effect of beneficial SNVs is estimated to be about 1.8 (1.5-2.2 HDI 95%).

#### Epistasis

We also estimate the epistasis between the beneficial SNVs. The epistasis parameter **η** is estimated to be about -0.3 (−1-0.443 HDI 95%), which implies sign epistasis and likely even reciprocal sign epistasis. To further validate this estimate, we used simulations to determine if the observation that pairs of beneficial SNVs rarely reside on the same genome (Fig. 2B) could be explained by a two-locus bi-allele Wright-Fisher model with just mutation, genetic drift, and strong selection, but without epistasis. Our simulation results strongly suggest such a model cannot explain the experimental results (Fig. S7), providing further support for sign epistasis between beneficial SNVs in MS2.

## Discussion

Both selection and sequencing errors can complicate the estimation of mutation rates from sequencing data (Gelbart et al., 2020; Peck & Lauring, 2018). Nevertheless, we show that our approach, which applies Bayesian inference to evolutionary experimental data using simulations and artificial neural networks, performs well on synthetic simulated data, and allows us for the first time to estimate the mutation rate of MS2 with a narrow confidence interval. In contrast to methods based on single-locus models, we set out to develop a multi-locus model that can account for background selection on full viral genomes as well as for standing variation in the founding population. This represents a challenge: even with the short genome of the MS2 virus, a high mutation rate leads to high genetic diversity (Fig. 3D), which makes the number of possible haplotypes huge. Thus, our model groups genomes by the number of mutant alleles they accumulated in several mutation types (Fig. 1). This model is simple enough to simulate efficiently but complex enough to capture the genome frequency dynamics. Furthermore, SNPE is also computationally more efficient than sampling methods such as REJ-ABC due to its application of neural density estimators as well as amortization (Avecilla et al., 2021).

Our estimate of the MS2 mutation rate is roughly *U=*0.2 mutations per genome per replication cycle (0.056-0.73 HDI 95%; Table 2). Reassuringly, we find that our method is robust to sequencing errors, which present a challenge for any method that relies on rare SNV frequencies (Gelbart et al., 2020, Acevedo et al., 2014). Compared with a close relative of MS2, qβ, our inferred mutation rate is an order-of-magnitude lower than the early estimates of the mutation rate of qβ, which were around *U=6*.*5* (Batschelet et al., 1976; Domingo et al., 1976). These studies quantified the mutation rate of one specific nucleotide at position 40 of the genome of qβ. Therefore, their estimates may be biased for that specific site, while our estimates represent a genome-wide mutation rate. A more recent estimate of the Qβ mutation rate at about *U=0*.*08* was based on phenotypic scores (García-Villada & Drake, 2012). Another study (Bradwell et al., 2013) corrected this estimate using the distribution of fitness effects obtained previously for this phage (Domingo-Calap et al., 2009) and estimated a mutation rate of *U=0*.*6*. This estimate is three times higher than our estimate, and within our estimated HDI. Importantly, the difference between mutation rate estimates for Qβ and MS2 may represent true biological differences between the two phages; indeed, their genomes share only ∼45% identity.

The inferred mutation rate, together with the very large population size (*θ=N·U/3,500=1,142*), puts this virus population in the regime of rapid adaptation unlimited by mutation (Feder et al., 2021). In this regime, soft sweeps are expected to occur and leave their mark on the genome. Indeed, we find evidence for multiple beneficial SNVs segregating in the population in multiple genotypes (Fig. 2). It would be interesting to estimate *θ* from SNV polymorphism dynamics (Feder et al., 2021; Pennings & Hermisson, 2006) to see if it agrees with our inference results, which directly estimates the mutation rate, as the population size is known in our experiment.

The posterior distribution of the non-beneficial nonsynonymous fitness effect *w*_*ns*_ is wide (Table S2). On the one hand, it is reassuring that there is a clear difference from the posterior of the non-beneficial synonymous fitness effect *w*_*s*_ (Fig. 3). However, the MAP estimate of 0.7 for *w*_*ns*_ is higher than previous estimates in the literature that found high frequency of lethal SNVs (Cuevas et al., 2012; Sanjuán et al., 2004). This difference may be explained by our use of a single fitness effect parameter to describe for all non-beneficial nonsynonymous SNVs, the distribution of which is likely bi-modal, with one ‘peak’ for lethal mutations and one ‘peak’ for slightly deleterious mutations (Cuevas et al., 2012; Domingo-Calap et al., 2009; Sanjuán, 2010; Sanjuán et al., 2004).

Epistatic effects are notoriously difficult to estimate from empirical data as they require departing from simple linear models, and more complex models require more data and stronger effect sizes to achieve statistical power (Elena et al., 2010). We first suspected epistatic effects when we observed that the beneficial SNVs are rarely found together on the same genome (Fig. 2B). To address this, we added an epistasis parameter to our model (Eq. 4). All our inferences estimated the epistasis parameter to be below 0.5, implying sign epistasis (Fig. 5B). Notably, this inference relied on the L-LR summary statistic, which uses labeled beneficial SNVs.

Interestingly, all four beneficial SNVs reside on the same RNA structure (Fig. 2D) and are close to the ribosomal binding site of the lysis protein. These SNVs are synonymous for the coat protein and three are non-synonymous for the lysis protein, whereas the fourth is not within the lysis gene. This may suggest that these beneficial SNVs either affect lysis translation, or change the lysis protein itself. Moreover, previous work has inferred that each of the single SNVs increases lysis expression, whereas a double mutant creates an unstable RNA structure that does not allow for lysis protein production and is therefore deleterious (Betancourt, 2009). Increased lysis expression in single mutants may allow the completion of two replication cycles in a single passage, which provides a strong fitness benefit. An alternative explanation for the beneficial effect of these SNVs is that these mutations decrease lysis expression, which gives the genome more time to replicate and create more progeny.

## Conclusions

Here we applied recent innovations in likelihood-free inference that use neural-network density estimators to directly approximate the posterior distribution rather than sampling from it, and long-read sequencing that covers the full length of a viral genome. This allowed us to efficiently and precisely infer the joint posterior distributions of the parameters governing the genome frequency dynamics in an evolutionary experiment with the MS2 bacteriophage, thereby estimating its mutation rate and fitness effects.

## Acknowledgements

We thank Grace Avecilla, David Gresham, Carmel Farage, and Uri Gophna for discussions. This work was supported in part by the Israel Science Foundation (ISF 552/19, YR), the US–Israel Binational Science Foundation (BSF 2021276, YR), Minerva Stiftung center for Lab Evolution (YR), by a European Research Council starting grant (852223 RNAVirFitness, AS), and by a fellowship from the Edmond J. Safra Center for Bioinformatics at Tel-Aviv University (NBN).

## Supplementary materials

**Supplementary Table 1.** Sequencing Coverage. Number of reads in each dataset.

| Replica | Passage | Sequencing Coverage |
| --- | --- | --- |
| A | 3 | 1343 |
|  | 7 | 4414 |
|  | 10 | 1552 |
| B | 3 | 2347 |
|  | 7 | 2604 |
|  | 10 | 1230 |
| C | 3 | 2446 |
|  | 7 | 1333 |
|  | 10 | 3350 |

**Supplementary Table 2.** Parameters estimates for ensemble SNPE.

| Parameter | Notation | Short-reads summary statistic |  | Long-reads summary statistic |  | Labeled long-reads summary statistic |  |
| --- | --- | --- | --- | --- | --- | --- | --- |
|  |  | MAP | HDI 95% | MAP | HDI 95% | MAP | HDI 95% |
| *Non-beneficial synonymous fitness effect* | $w_s$ | 0.935 | (0.451, 1) | 0.946 | (0.482, 1) | 0.897 | (0.651, 1) |
| *Non-beneficial nonsynonymous fitness effect* | $w_{ns}$ | 0.847 | (0.259, 0.98) | 0.797 | (0.256, 0.99) | 0.679 | (0.311, 0.922) |
| *Beneficial fitness effect* | $w_b$ | 2.012 | (1.334, 2.887) | 1.769 | (1.191, 2.582) | 1.822 | (1.508, 2.246) |
| *Beneficial synonymous SNV probability* | $p_{bs}$ | 0.0026 | (0, 0.006) | 0.001 | (0.001, 0.005) | 0.0008 | (0, 0.0022) |
| *beneficial nonsynonymous SNV probability* | $p_{bns}$ | 0.0072 | (0.003, 0.01) | 0.006 | (0.006, 0.01) | 0.0073 | (0.0032, 0.01) |
| *average number of synonymous SNVs per genotype in the initial population* | $M_s$ | 0.539 | (0.455, 0.6) | 0.545 | (0.545, 0.6) | 0.533 | (0.425, 0.6) |
| *average number of nonsynonymous SNVs per genotype in the initial population* | $M_{ns}$ | 0.773 | (0.705, 0.874) | 0.781 | (0.781, 0.879) | 0.799 | (0.711, 0.887) |
| *Initial log-fitness correlation* | $\delta$ | 0.049<br>9 | (0, 1.361) | 0.03 | (0, 1.242) | 0.0698 | (0,1.112) |
| *Epistasis* | $\eta$ | -0.1 | (-1, 1221) | 0.296 | (-1, 1.395) | -0.317 | (-1, 0.443) |

**Supplementary Figure S1.**
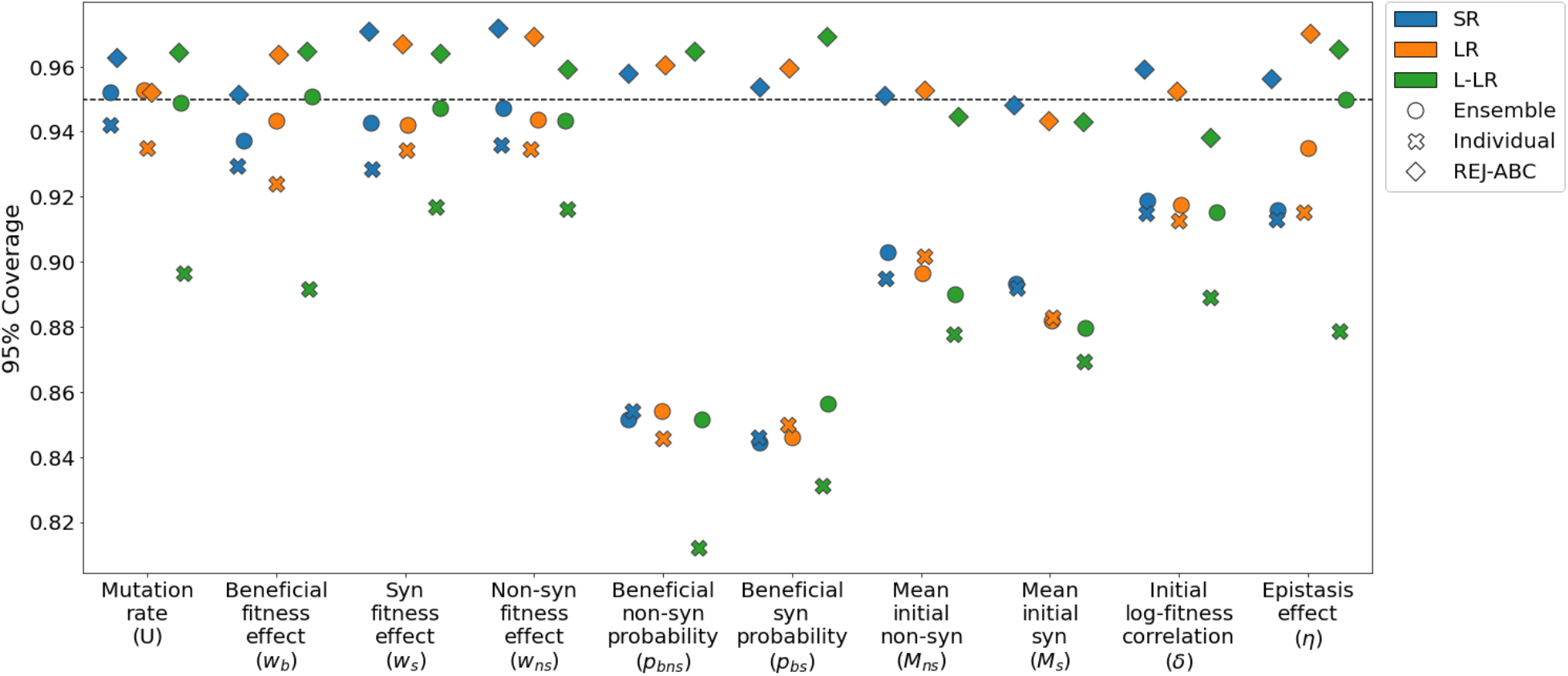
95% Coverage property of different inference methods on synthetic data. The coverage property tests that the inferred 95% credible interval contains the true parameter in 95% of the cases. Ensemble SNPE (eight estimators, each trained on 10,000 training examples) has better coverage than individual SNPE (trained on the entire 80,000 training dataset). REJ-ABC almost always has higher coverage, a result of much wider confidence intervals; REJ-ABC is also less accurate (Fig. 3). SR, LR, and L-LR stand for short-reads, long-reads and labeled long-reads summary statistic, respectively.

**Supplementary Figure S2.**
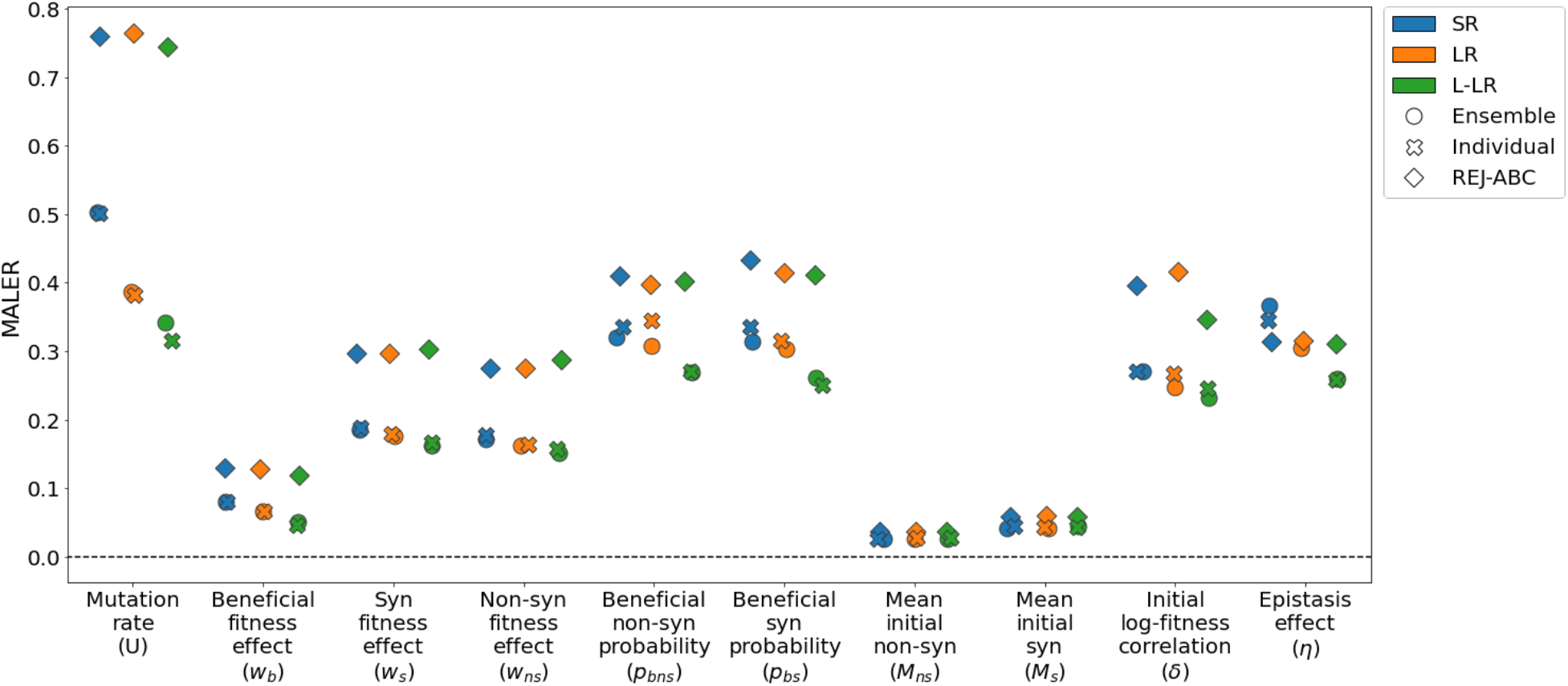
Estimation accuracy for different inference methods on synthetic data. The mean absolute log error ratio of the maximum a-posteriori (MAP) estimates and the true value. Lower is better, zero is best. Evaluation done on a synthetic dataset of 2,000 simulations that were simulated by using parameters sampled from the same prior as the training dataset. SR, LR, and L-LR stand for short-reads, long-reads and labeled long-reads summary statistic, respectively.

**Supplementary Figure S3.**
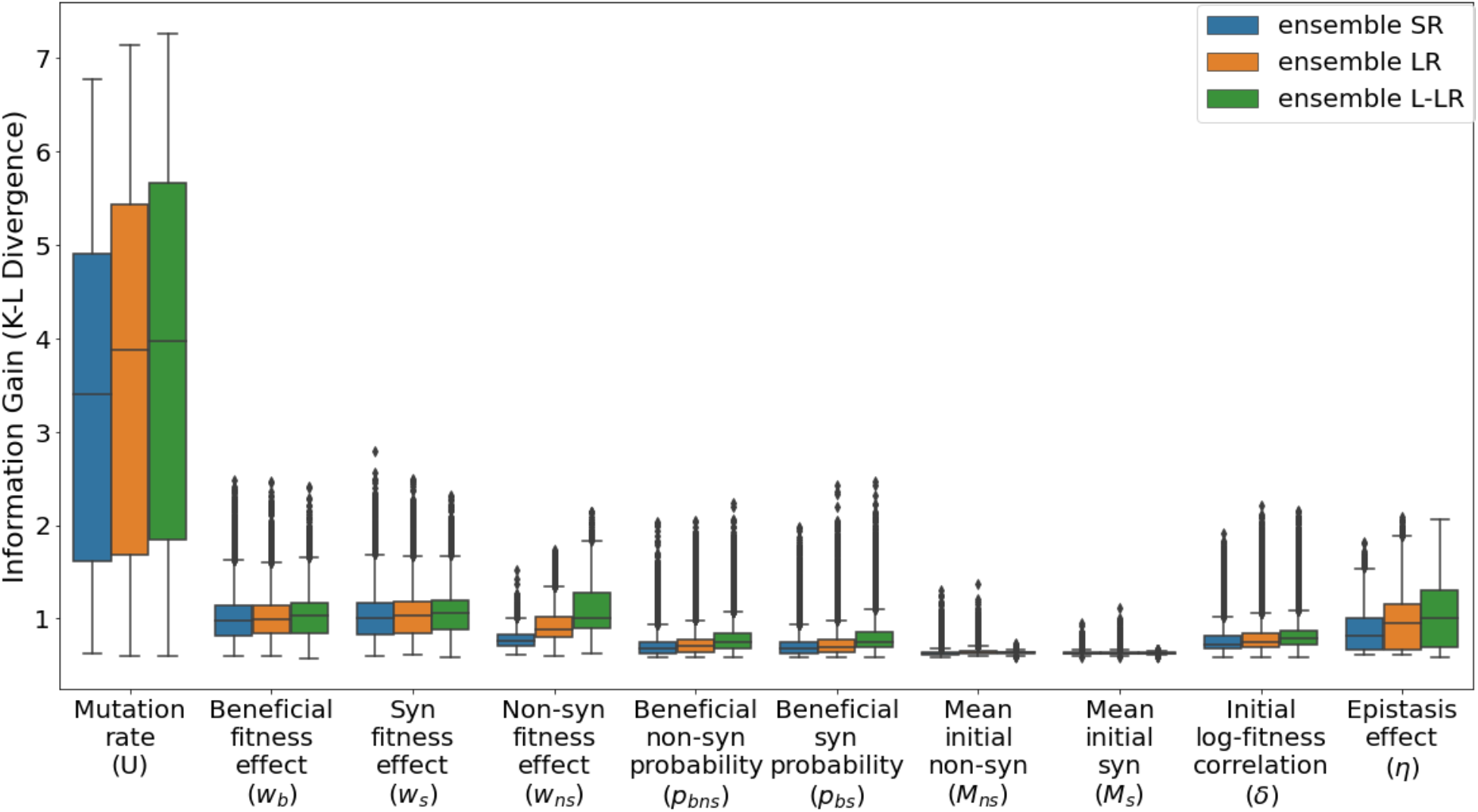
Information Gain for different summary statistics. The information gain is the Kullback-Leibler (KL) divergence between posterior and prior distributions. Higher is better. Evaluation done on a synthetic dataset of 2,000 simulations that were simulated by using parameters sampled from the same prior as the training dataset. SR, LR, and L-LR stand for short-reads, long-reads and labeled long-reads summary statistic, respectively.

**Supplementary Figure S4.**
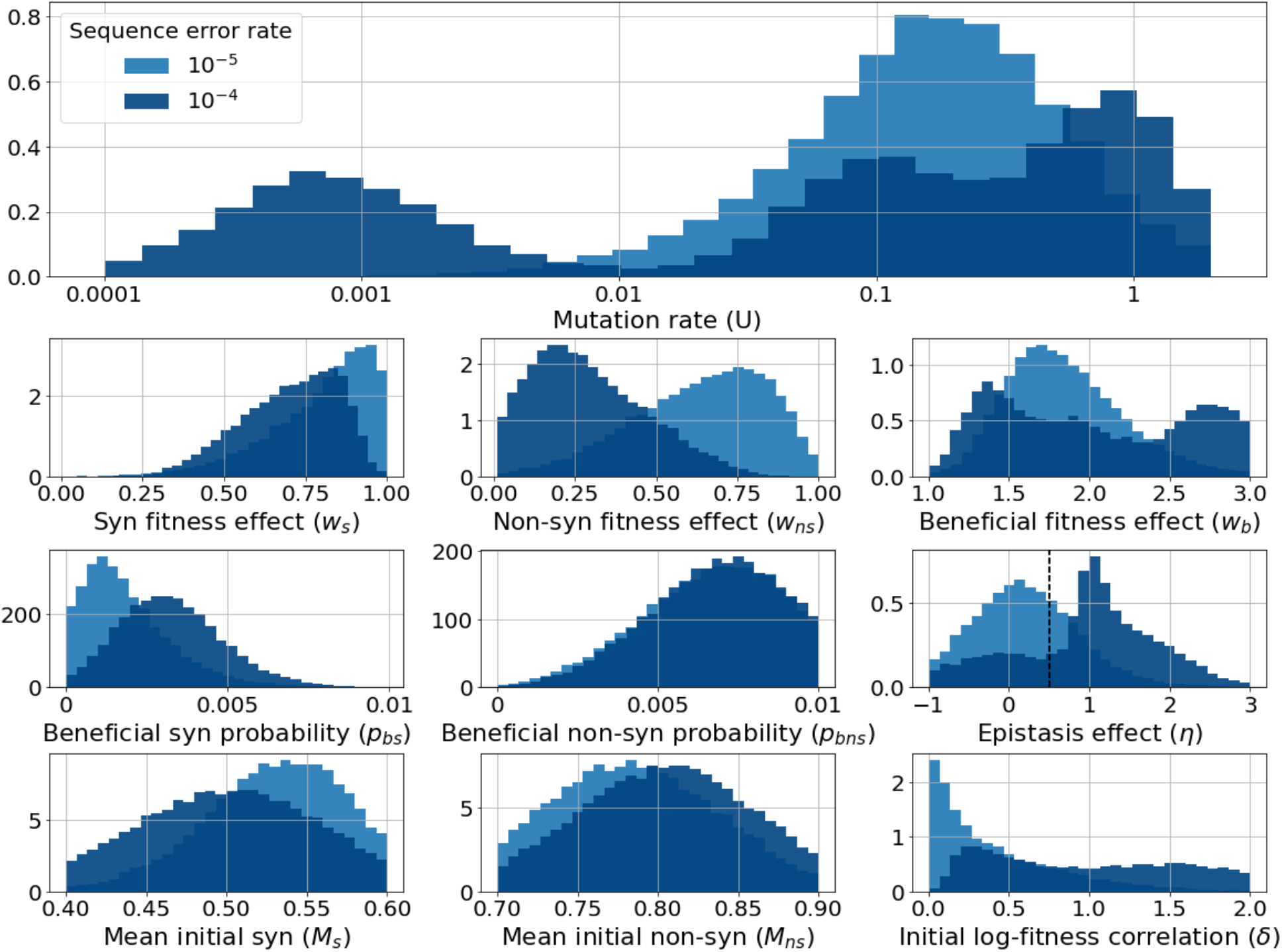
Effect of sequencing error on parameter estimation: LR summary statistic. Marginal posteriors of all model parameters compared with tenfold higher sequencing error rate. Inferences are from SNPE with long-reads (LR) summary statistic.

**Supplementary Figure S5.**
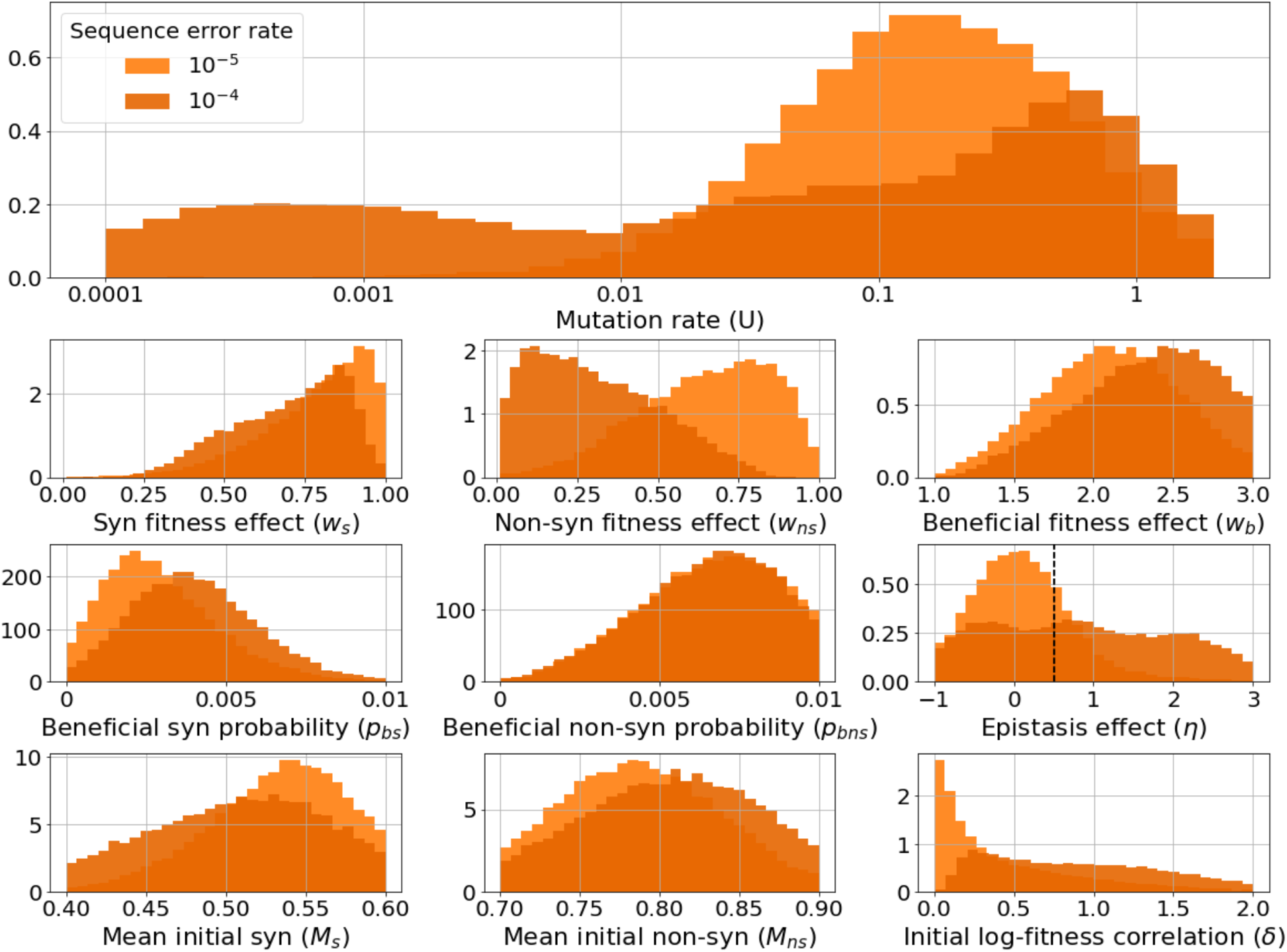
Effect of sequencing error on parameter estimation: SR summary statistic. Marginal posteriors of all model parameters compared with tenfold higher sequencing error rate. Inferences are from SNPE with short-reads summary statistic (SR).

**Supplementary Figure S6.**
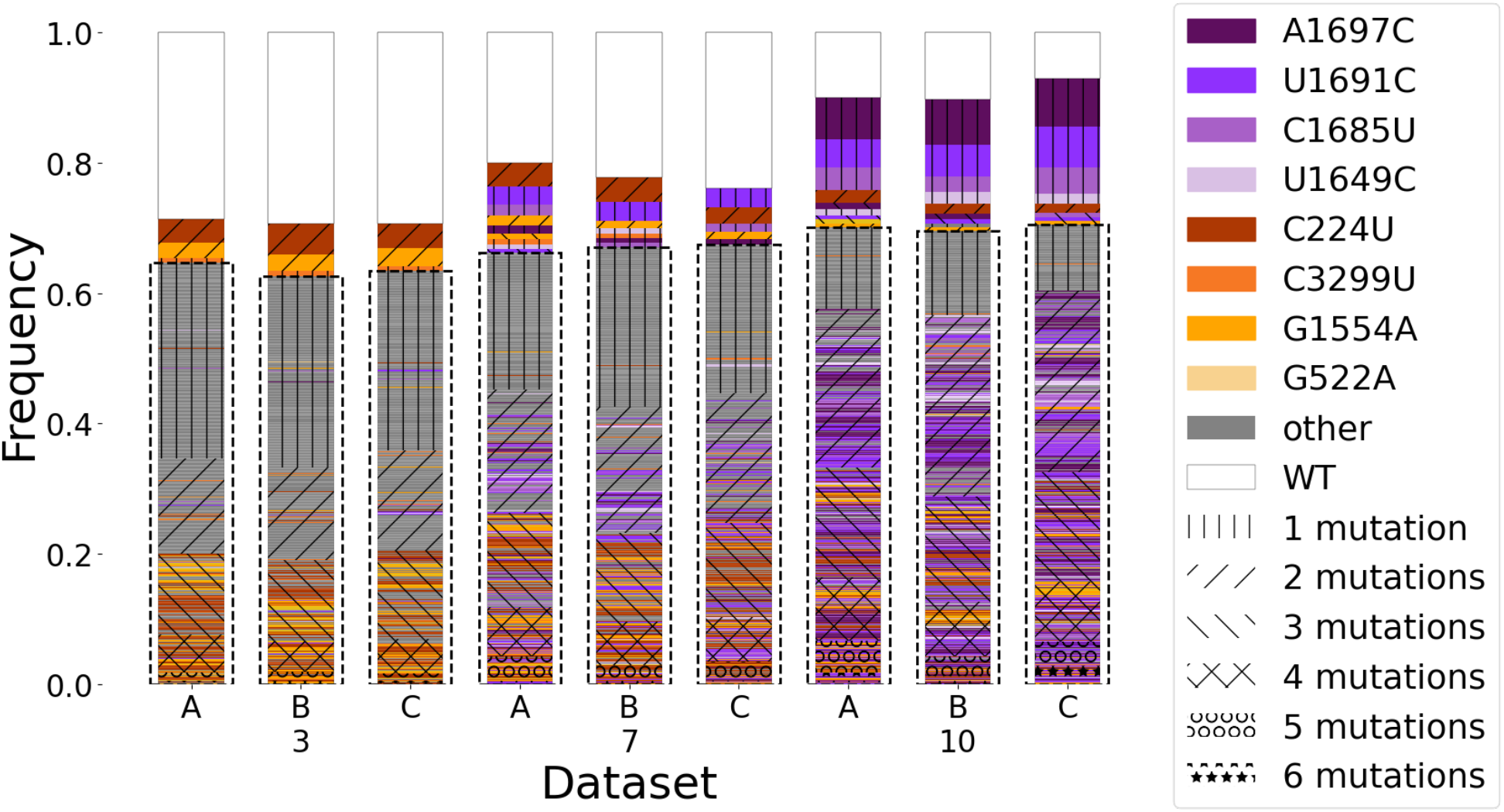
Genetic Diversity. For each replica and passage we show the different genotypes color coded by the SNVs they bear. When a genotype contains more than one high frequency SNV its color is defined by the first one in order of appearance in the legend. The markings on the bars represent the number of SNVs per genotype. The genotypes are ordered by their frequency in each dataset and the number of mutations within them. Rare genotypes are defined as present in less than 0.5% of the sample.

**Supplementary Figure S7.**
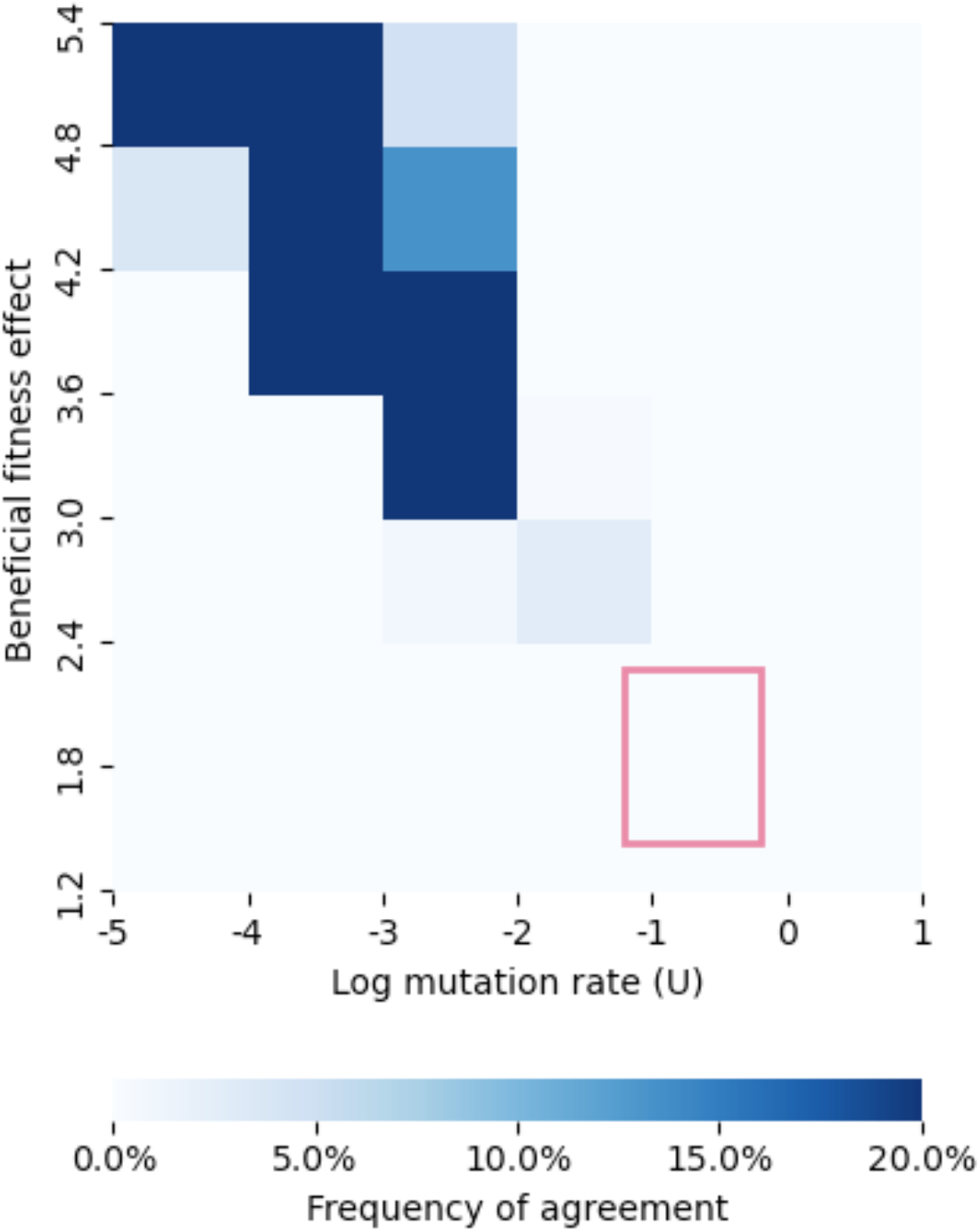
Testing whether mutation and selection alone allow negative linkage disequilibrium. We simulated 12,600,000 instances of a Wright-Fisher two-locus bi-allelic model with selection, mutation, and drift with mutation rate *U* sampled uniformly from (5 · 10^−5^, 10), fitness effects sampled uniformly from (1.2, 5.4), and effective population size of 2·10^7^ for 10 generations. For each simulation we determined if, at passage 10, the double mutant frequency was lower than or equal to the empirical frequency (Fig. 2) and if the single mutant frequency was higher than or equal to the empirical frequency. The figure shows the frequency of such agreements between simulations and empirical data. Such agreements are likely for extremely low mutation rates (dark blue), which are uncharacteristic for viruses, and unrealistically high fitness effect values. The red box marks the boundaries of the HDI 95% posterior distribution from our main analysis for U and *w*_*k*_, and 0% agreement was found in this scenario.

**Figure S8.**
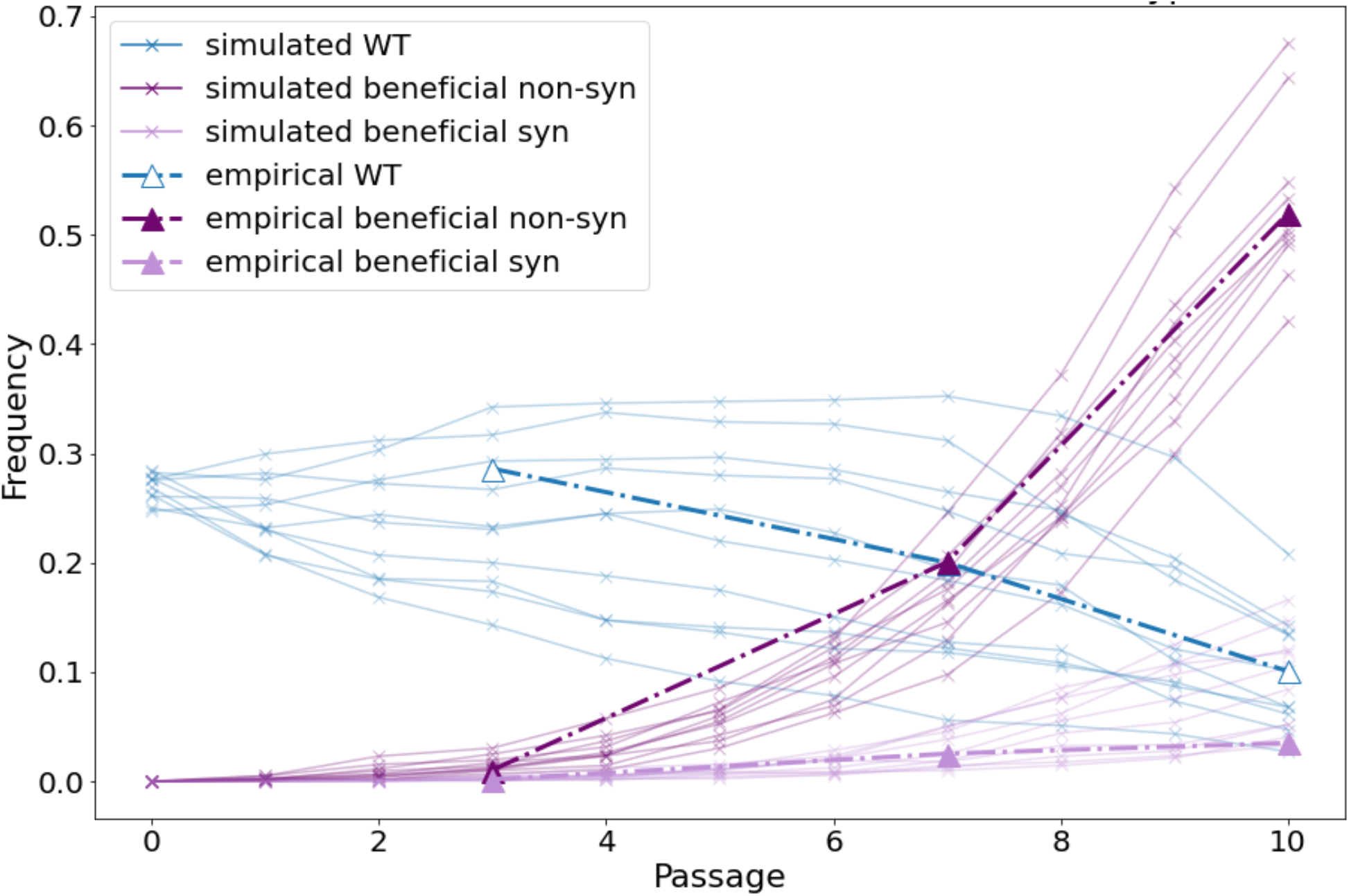
Posterior predictive check of wildtype and beneficial SNVs frequency dynamics. The figure compares 10 posterior predictions of the frequency dynamics (solid lines) to the empirical data of population A (dashed bold lines). Predictions were generated by simulating the evolutionary model with parameter sets sampled from the posterior distribution inferred with ensemble SNPE with the L-LR summary statistic.

**Supplementary Figure S9.**
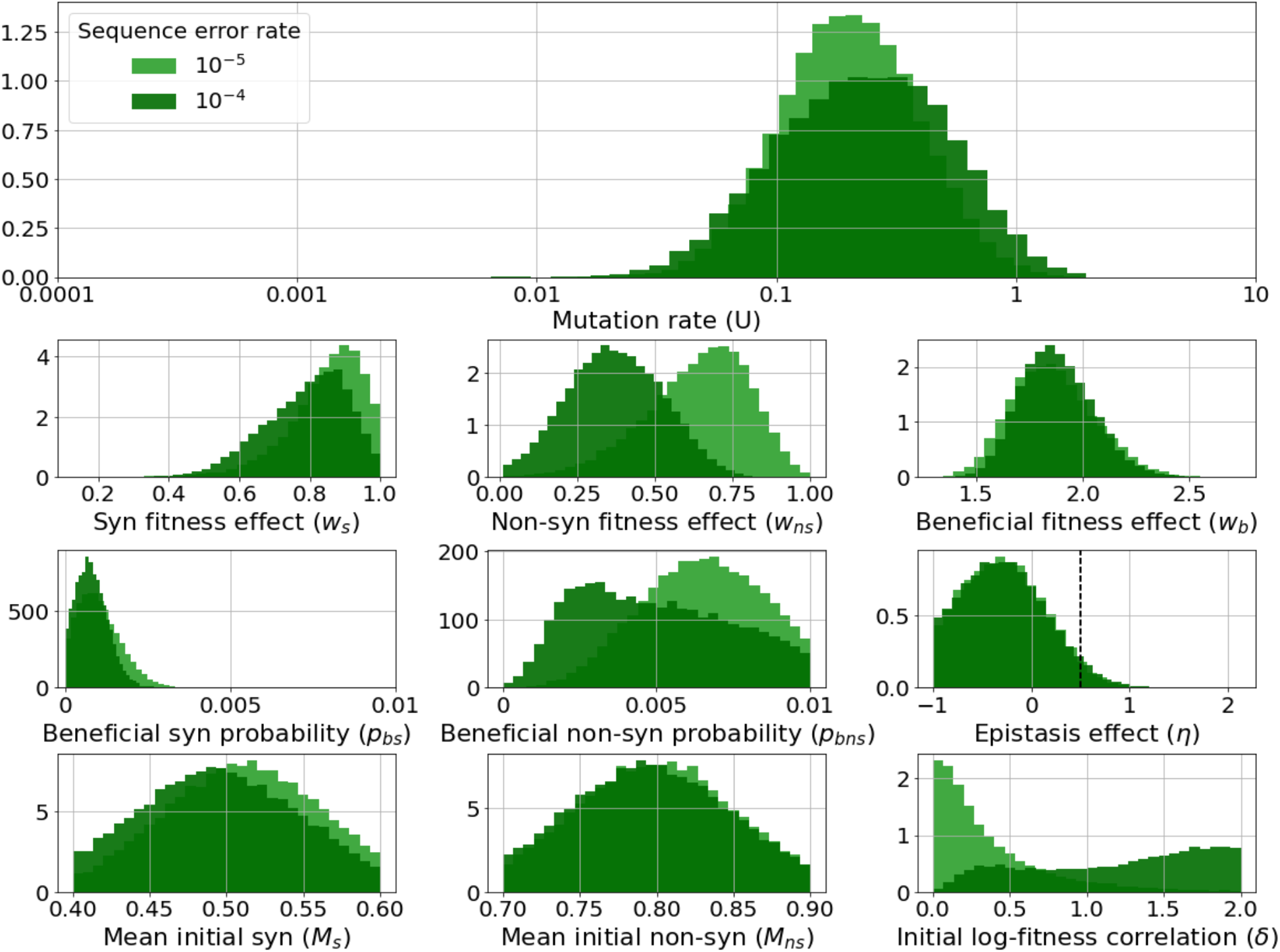
Effect of sequencing error on parameter estimation: L-LR summary statistic. Marginal posteriors of all model parameters compared with tenfold higher sequencing error rate. Inferences are from SNPE with labeled long-reads summary statistic (L-LR).

## Notes

https://github.com/Stern-Lab/ms2-mutation-rate

https://www.ncbi.nlm.nih.gov/sra/PRJNA902661

https://zenodo.org/record/7486851

### Competing Interest Statement

The authors have declared no competing interest.

